# Comparative analysis of human brain organoids of brainstem and midbrain at single-cell resolution

**DOI:** 10.1101/2020.09.02.279380

**Authors:** Kaoru Kinugawa, Joachim Luginbühl, Takeshi K. Matsui, Nobuyuki Eura, Yoshihiko M. Sakaguchi, Jay W Shin, Kazuma Sugie, Jens C. Schwamborn, Eiichiro Mori

## Abstract

Human brain organoids provide us the means to investigate human brain development and neurological diseases, and single-cell RNA-sequencing (scRNA-seq) technologies allow us to identify homologous cell types and the molecular heterogeneity between individual cells. Previously established human brain organoids of brainstem (hBSOs) and midbrain (hMBOs) were analyzed by scRNA-seq, but the difference in cellular composition between these organoids remains unclear. Here, we integrated and compared the single-cell transcriptome of hBSOs and hMBOs. Our analysis demonstrated that the hBSOs and hMBOs contain some unique cell types, including inflammatory and mesenchymal cells. Further comparison of the hBSOs and hMBOs with publicly available scRNA-seq dataset of human fetal midbrain (hMB) showed high similarity in their neuronal components. These results provide new insights into human brain organoid technologies.

## INTRODUCTION

Recenly developed three-dimensional *in vitro* human organ models in-a-dish, organoids, are derived from human pluripotent stem cells, such as embryonic stem cells (ESCs) (1) and induced pluripotent stem cells (iPSCs) (2,3). They mimic the composition of human organs, including the brain (4,5), colon (6) and kidney (7). They have been used to elucidate the pathology of central nervous system disorders (4,8). Various methods for region-specific human brain organoids have been developed, such as hypothalamic (8), midbrain (8-11) and cerebellar organoids (12). Previously, we designed and constructed new methods for generating human brainstem organoids (hBSOs) (13) and human midbrain organoids (hMBOs) (11). These methods are expected to be useful tools for *in vitro* drug discovery and neurodegenerative disease modeling, such as Parkinson’s disease (PD) and DiGeorge syndrome (14).

In our previous studies, we performed single-cell RNA-sequencing (scRNA-seq) on hBSOs and hMBOs. The scRNA-seq technologies allow us to analyze transcriptomes of a variety of samples at single-cell resolution (15-18). Rapid development of scRNA-seq technologies has provided us valuable insights into complex biological process. The scRNA-seq analysis demonstrated that hBSOs and hMBOs contained various cell types and neuronal subtypes, including dopaminergic, glutamatergic and GABAergic (13,19). However, the difference in transcriptome between hBSOs and hMBOs remained unclear. In this study, we compared the single-cell transcriptome of hBSOs and hMBOs. These organoids contain some unique cell types, such as mesodermal-derived cells. Further comparative analysis demonstrated the similarity between these organoids and fetal midbrain. These results give us new insights about human brain organoids.

## MATERIALS AND METHODS

### Clustering analysis

The two scRNA-seq datasets that we previously reported were analyzed using Seurat v.3.0.0 R package (20). We filtered the cells and genes as described in previous studies (13,19). We collected a total of 3,640 cells (2,345 cells at hBSOs and 1,295 cells at hMBOs). The two datasets were log-normalized and highly variable genes were identified using the variance-stabilizing transformation method. The two datasets were integrated using canonical correlation analysis (CCA) method, “FindIntegrationAnchors” and “IntegrateData” functions in Seura (21). Principal components analysis (PCA) was performed on the integrated datasets. Based on the top 30 principal components (PCs), clustering was performed using the shared nearest neighbor (SNN) modularity optimization with resolution set to 0.8. All cells were classified into 12 clusters. Each cluster was identified based on differentially expressed genes using the “FindAllMarkers” function in Seurat (Table S1).

We analyzed a publicly available human midbrain (hMB) scRNA-seq dataset (GSE76381) generated by Manno et al (22). The hMB datasets contained a total of 1,977 cells from human embryos. We collected a total of 5,617 cells (1,977 cells at hMB, 2,345 cells at hBSOs and 1,295 cells at hMBOs). The datasets of hMB and the two organoids were integrated using CCA method, “FindIntegrationAnchors” and “IntegrateData” functions. After PCA, clustering was performed based on the top 30 PCs using SNN modularity optimization with a resolution of 0.8. All cells were classified into 14 clusters. Cluster identities were determined by cluster gene markers using the “FindAllMarkers” function in Seurat (Table S2).

### Pearson correlation coefficient analysis

All 5,617 cells were classified into 14 clusters. In order to perform our correlation analysis, we combined some clusters and assigned these clusters new identities. We assigned “Ex/Inhi N1” and “Ex/Inhi N2” to “Ex/Inhi N”, as well as “NP1” and “NP2” to “NP”. We obtained a total of 12 different cell type components. The hBSOs, hMBOs and hMB contained various components in different cell numbers (Table S3). Cell components that contained less than 5 cells (e.g. FB Prog of hMB, OPC of hBSO) were excluded from subsequent analysis. Using “AverageExpression” function in Seurat, we computed average gene expression level for each cluster between hBSOs, hMBOs and hMB. Using “CellScatter” function in Seurat, we computed Pearson correlation coefficient between each cell component (Table S4, S5, S6).

### Differentially expressed gene analysis

We detected genes that were differentially expressed between one cluster and any other cluster. We applied the “FindAllMarkers” function in Seurat using default parameters.

## RESULTS

### hBSOs and hMBOs contained divergent composition

Previously, we performed scRNA-seq on hBSOs and hMBOs and showed that these two organoids contained various cell types such as neuronal progenitor, neuron and radial glia (13,19). The scRNA-seq analysis supported the finding that hBSOs and hMBOs recapitulate the human brainstem and midbrain, respectively (Figure 1A). To better understand the difference between hBSOs and hMBOs, we performed an integrative analysis on scRNA-seq datasets of the two organoids. After filtering out the cells by quality control, a total of 3,640 cells (2,345 cells at hBSOs and 1,295 cells at hMBOs) were divided into 12 cell clusters and projected into a two-dimensional (2D) space by Uniform Manifold Approximation and Projection (UMAP) (Figure 1B). We selected cluster-enriched differentially expressed genes (Table S1). Each cluster expressed marker genes for each cell type (Figure 1C, Figure S1) and were identified based on the gene expression profiles (Figure 2A). These results suggest that a total of 3640 cells were successfully classified into cell type-specific clusters.

**Figure 1.**
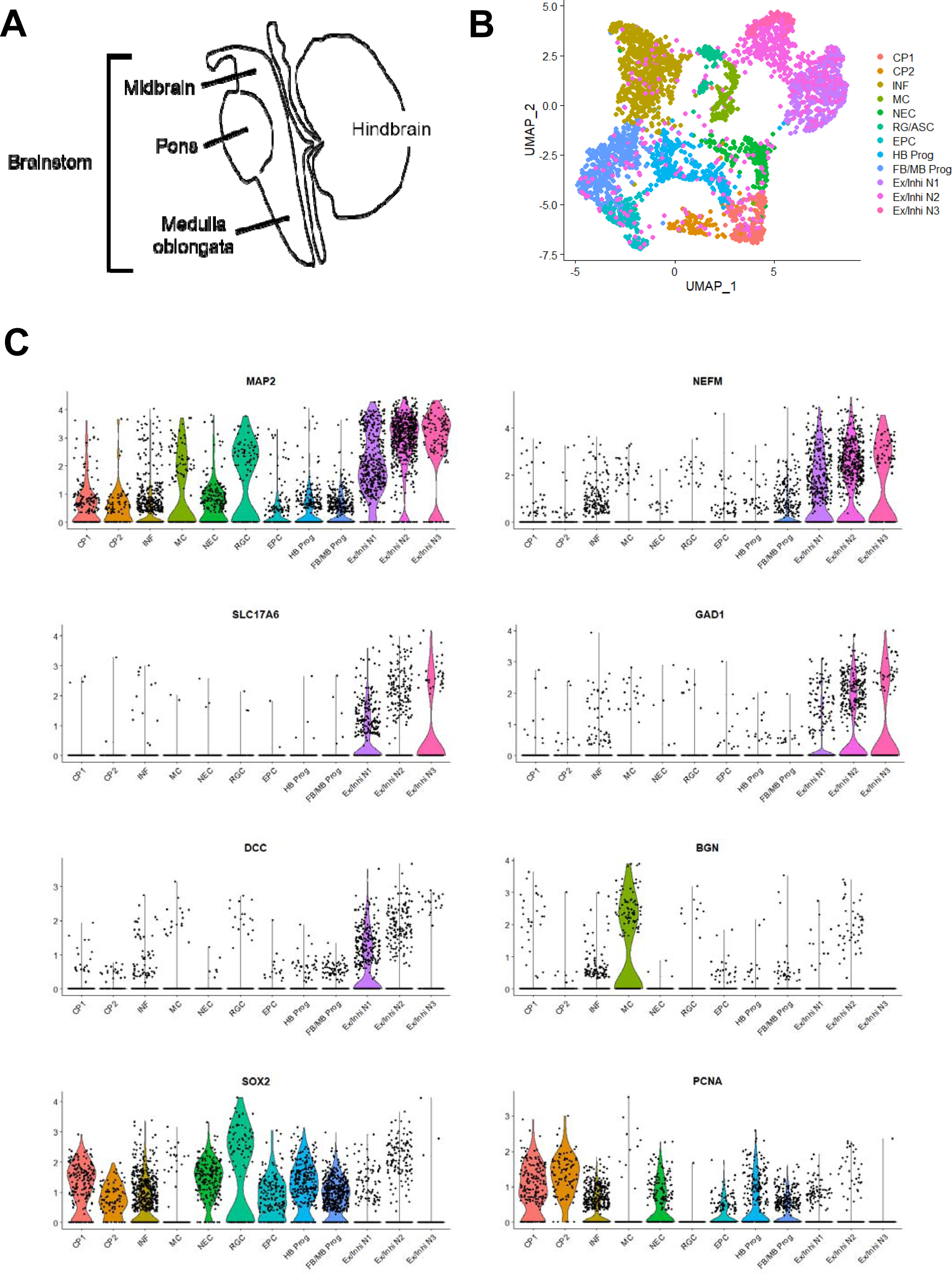
A total of 3640 cells from hBSOs and hMBOs were classified into 12 cell type-specific clusters. (A) Schematic drawings of the brainstem and hindbrain. (B) UMAP plot of total 3640 cells distinguished by cell types. Each dot corresponds to a single cell. CP: cycling progenitors; INF: inflammation; MC: mesencymal cells; NEC: neuroepithelial cells; RG/ASC: radial glia and astrocyte; EPC: ependymal cells; HB Prog: hindbrain progenitors; FB/MB Prog: forebrain and midbrain progenitors; Ex/Inhi N: excitatory and inhibitory neuron. (C) Violin plots show distribution of cell type-specific marker genes for 12 cell clusters.

**Figure 2.**
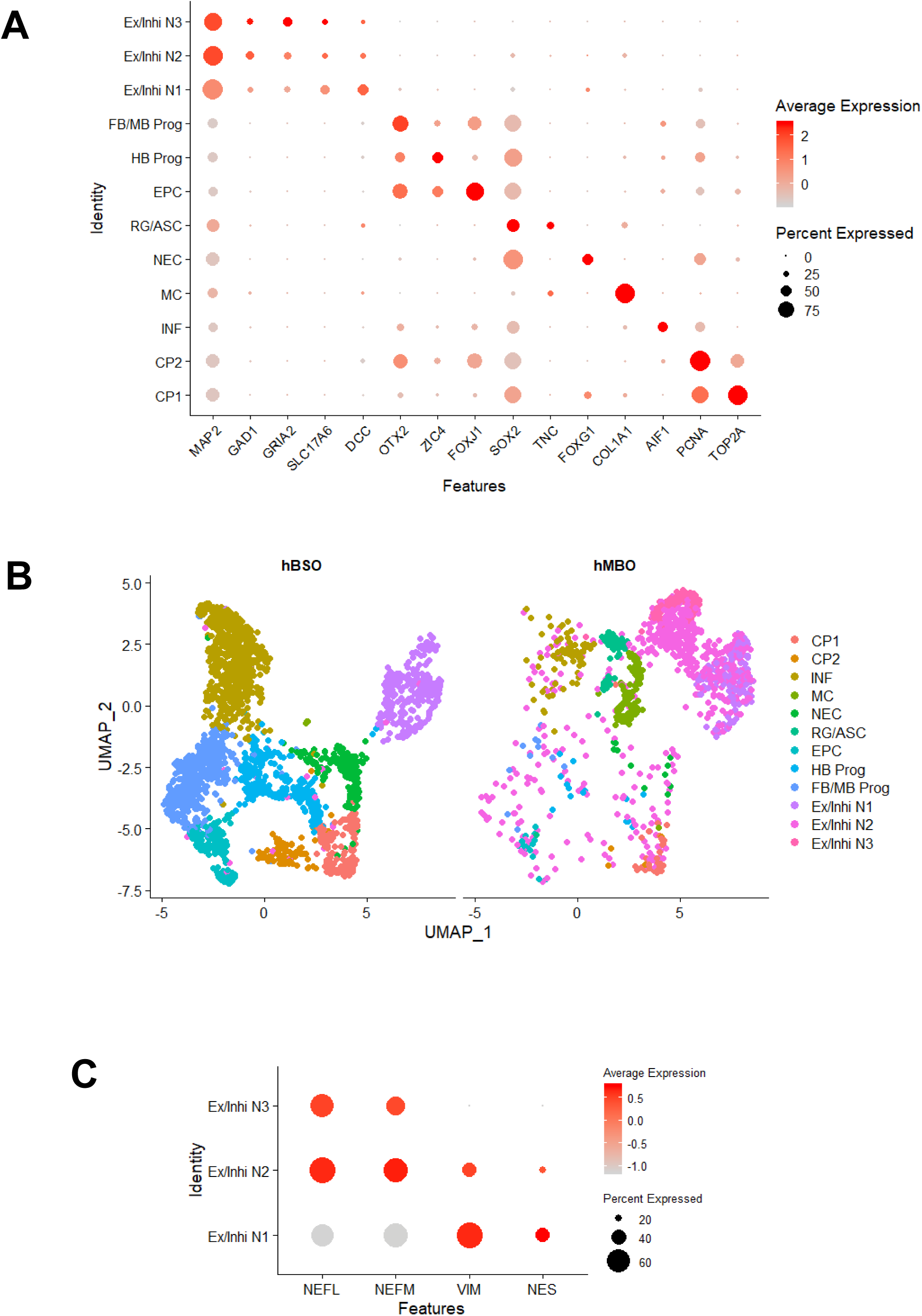
The hBSOs and hMBOs contained unique cell populations. (A) Dot plot shows the expression of cell type-specific marker genes for 12 cell clusters. (B) UMAP plot shows that hBSOs and hMBOs contain different cell clusters. (C) Dot plot shows the gene expression of intermediate filament markers for Ex/Inhi N1, Ex/Inhi N2, and Ex/Inhi N3 clusters.

HBSOs contained some clusters that were different from that of hMBOs (Figure 2B). For example, hBSOs contained the hindbrain progenitors (HB Prog) cluster, which expresses genes of cerebellar or medulla formation (*ZIC1, ZIC4*) (23). HBSOs also contained an inflammation (INF) cluster, which expresses the gene related to microglia function (*AIF1*). On the other hand, hMBOs contained different clusters (Figure 2B), such as the radial glia and astrocyte (RG/ASC) cluster. The RG/ASC cluster expressed *SOX2* and *TNC*, which drives astrocyte differentiation. HMBOs also contained the mesenchymal cells (MC) cluster. The MC cluster expressed genes of collagen synthesis, *COL1A1* and *BGN* (24). These results suggest that hBSOs and hMBOs each contain some unique cell populations.

HMBOs contained three excitatory and inhibitory neuron clusters (Ex/Inhi N1, Ex/Inhi N2, Ex/Inhi N3; Figure 2B). These clusters expressed the gene of different neuronal subtypes, including dopaminergic (*DDC*), GABAergic (*GAD1*), and glutamatergic (*GRIA2, SLC17A6*) neuronal markers (Figure S1). The hBSOs contained only Ex/Inhi N1 cluster, which highly expressed *VIM* and *NES*, intermediate filament markers for mesenchymal progenitor cells and neural stem cells (Figure 2C) (25). On the other hand, the Ex/Inhi N2 and Ex/Inhi N3 clusters highly expressed *NEFL* and *NEFM*, which are intermediate filament markers for differentiated neural cells (Figure 2C). These results suggest that hMBOs contain neuronal cells with different developmental stages.

### hBSOs and hMBOs contained similar composition compared to hMB

To compare our two organoids with human tissue, we used the scRNA-seq dataset from human fetal midbrain (22). A total of 5,617 cells (1,977 cells at hMB, 2,345 cells at hBSOs and 1,295 cells at hMBOs) were integrated and divided into 14 cell clusters (Figure 3A). Each cluster expressed marker genes for each cell type (Figure 3B, Figure S2) and was identified based on the gene expression profiles (Figure 4A). These results demonstrate that the 5,617 cells derived from three datasets (hMB, hMBOs and hBSOs) could be successfully classified into cell type-specific clusters.

**Figure 3.**
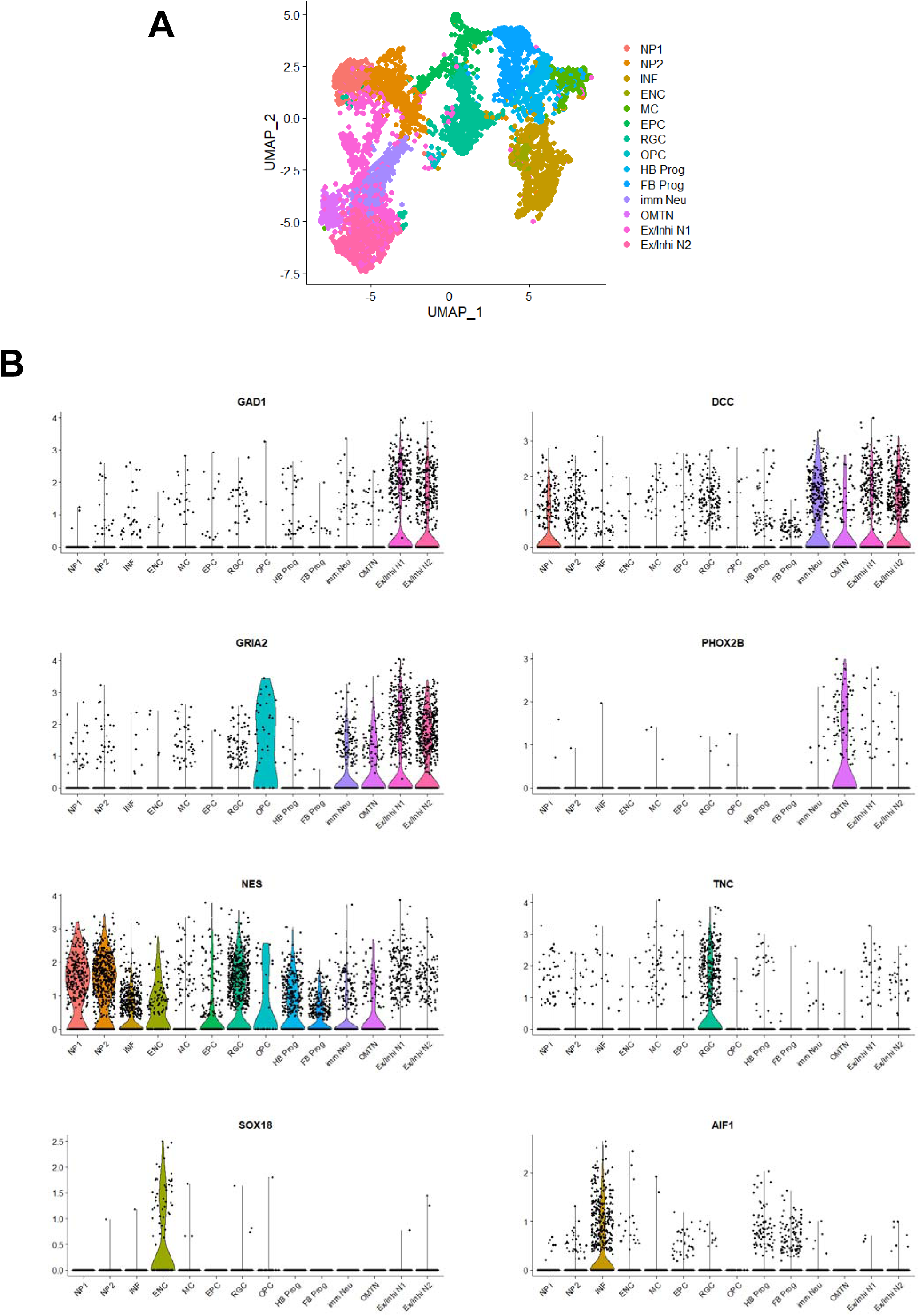
A total of 5617 cells from hMB, hBSOs and hMBOs were classified into 14 cell type-specific clusters. (A) UMAP plot of all 5617 cells distinguished by cell types. Each dot corresponds to a single cell. NP: neuronal progenitors; INF: inflammation; ENC: endothelial cells; MC: mesencymal cells; EPC: ependymal cells; RGC: radial glia cells; OPC: oligodendrocyte precursor cells; HB Prog: hindbrain progenitors; FB Prog: forebrain progenitors; imm Neu: immature neurons; OMTN: oculomotor and trochlear nucleus; Ex/Inhi N: excitatory and inhibitory neuron. (B) Violin plots show distribution of cell type-specific marker genes for 14 cell clusters.

**Figure 4.**
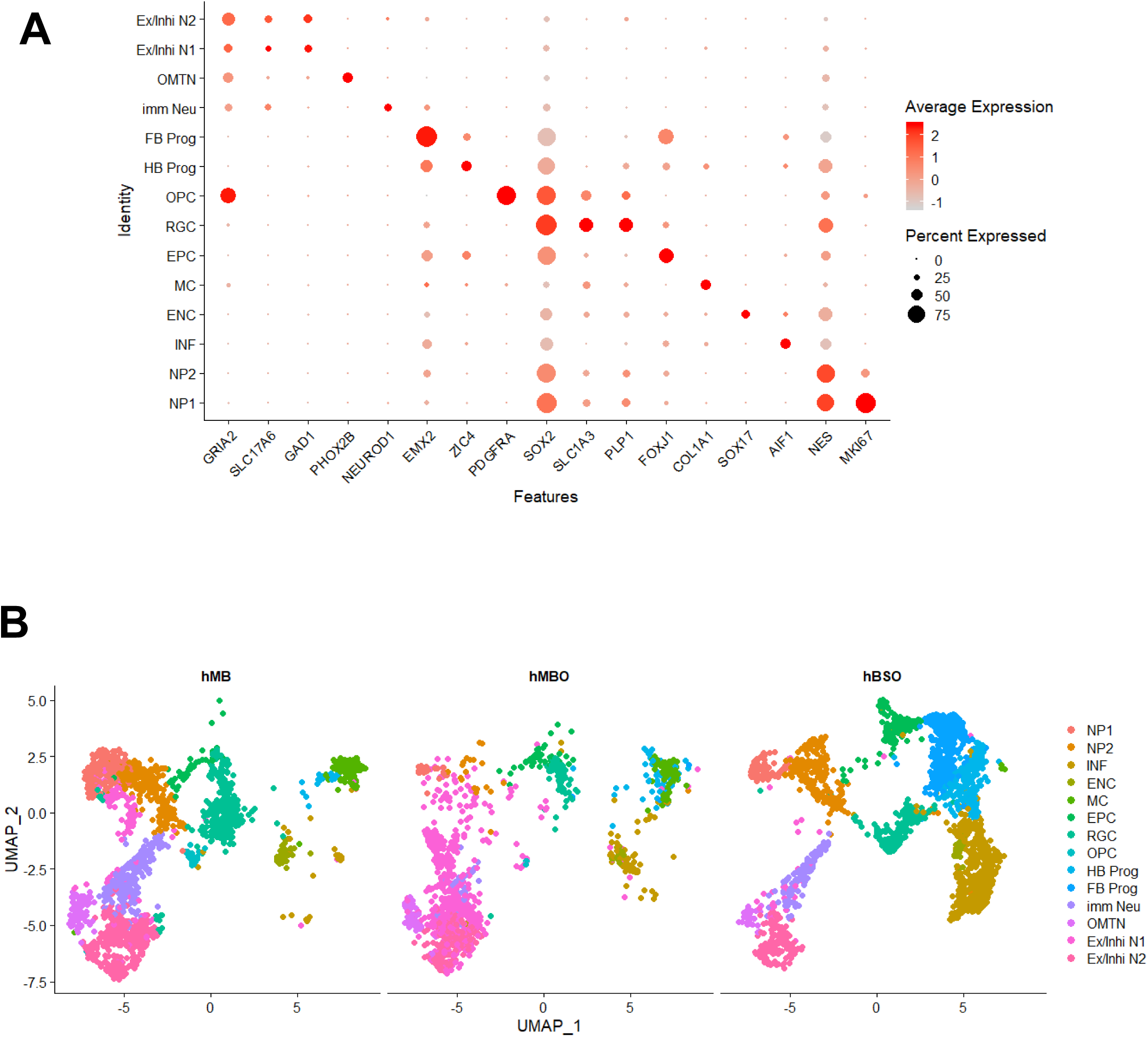
The hMB, hMBOs and hBSOs contained similar composition. (A) Dot plot shows the expression of cell type-specific marker genes for 12 cell clusters. (B) UMAP plot shows that the hMB, and hMBOs contain similar cell clusters.

The cell distribution pattern among the three datasets is partially similar to one another (Figure 4B). We also observed that, similar to hMB, the two organoids contained various neuronal lineage clusters like the neuronal progenitors (NP1, NP2), excitatory and inhibitory neurons (Ex/Inhi N1, Ex/Inhi N2), and oculomotor and trochlear nucleus (OMTN) clusters (Figure 4B). On the other hand, the two organoids did not contain oligodendrocyte precursor cells (OPC) cluster, which expressed *PDGFRA* and *OLIG1/2* (Figure 4B, Figure S2). These results suggest that the two organoids contain the similar neuronal components as that of hMB.

### hBSOs and hMBOs are similar to hMB based on correlation coefficient analysis

To assess the similarity between the two organoids and hMB, we calculated Pearson correlation coefficient (PCC) scores based on their transcriptional expression profiles (please refer to our Methods for more details). The PCC score between hBSOs and hMBOs was 0.63 (Figure 5A). Both hBSOs and hMBOs showed high correlations with hMBs, with PCC scores of 0.74 and 0.72, respectively (Figure 5A). To further investigate their transcriptional similarity, we calculated the PCC between every clusters that originated from hMB, hBSOs and hMBOs. For example, we observed that the Ex/Inhi N component of hMBOs showed high similarity with Ex/Inhi N of hMB (PCC, 0.77; Figure 5B, Table S4). NP component of hBSOs showed high similarity with NP of hMB (Figure 5C, Table S5). The neuronal components, Ex/Inhi N, OMTN and immature neurons (Imm Neu), showed high similarity among the three datasets (Figure 5B, 5C, 5D). These results further suggest that the two organoids share similar neuronal components with human fetal midbrain.

**Figure 5.**
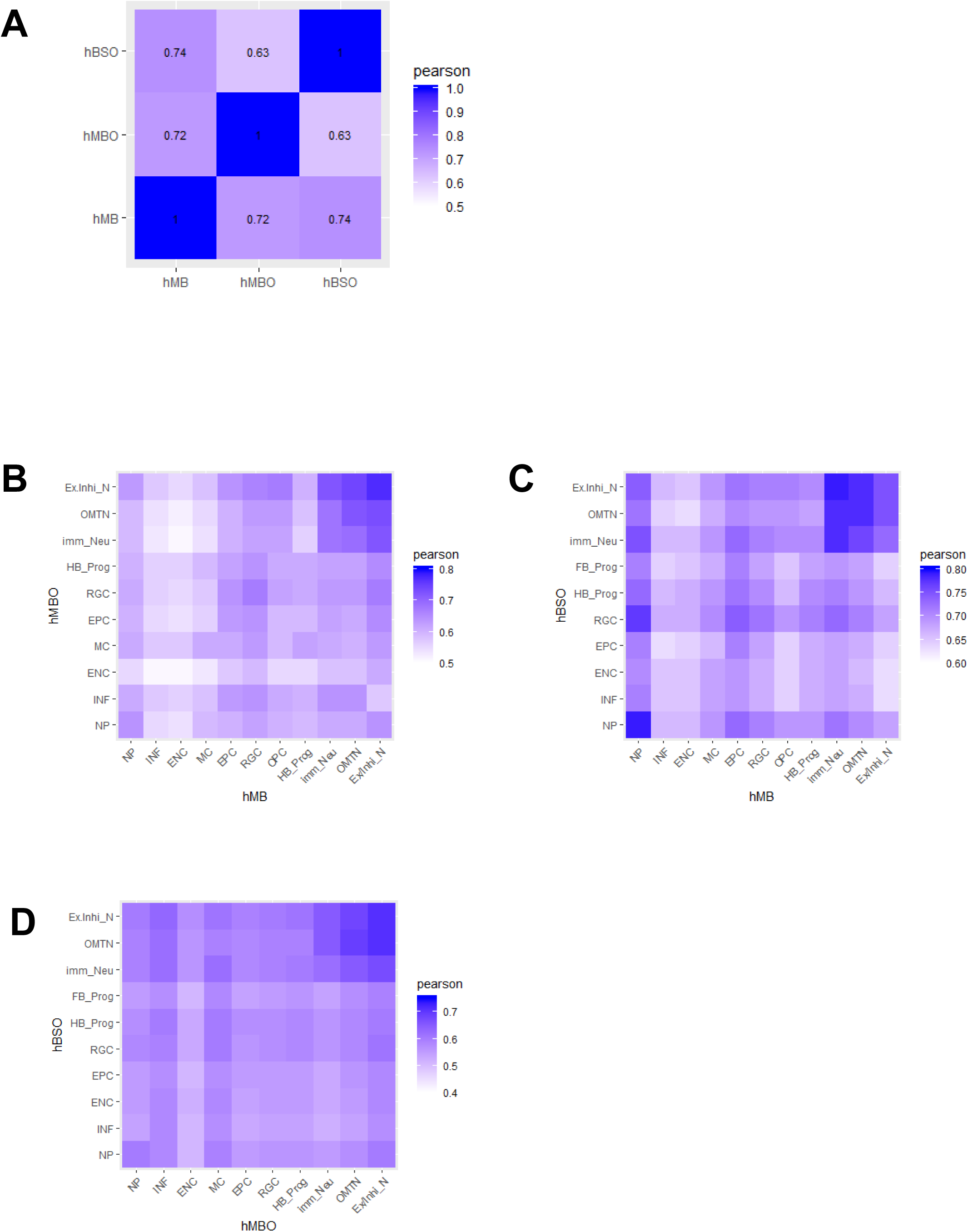
Pearson correlation coefficient (PCC) analysis shows similarity between hMB, hMBOs and hBSOs. (A) The PCC between hMB, hMBOs and hBSOs. (B) The PCC between hMBOs and hMB cell types. The heatmap shows correlation based on gene expression in each cell type. NP: neuronal progenitors; INF: inflammation; ENC: endothelial cells; MC: mesencymal cells; EPC: ependymal cells; RGC: radial glia cells; OPC: oligodendrocyte precursor cells; HB Prog: hindbrain progenitors; FB Prog: forebrain progenitors; imm Neu: immature neurons; OMTN: oculomotor and trochlear nucleus; Ex/Inhi N: excitatory and inhibitory neuron. (C) The PCC between hBSOs and hMB cell types. The heatmap shows correlation based on gene expression in each cell type. (D) The PCC between hBSOs cell types and hMBOs cell types. The heatmap shows correlation based on gene expression in each cell type.

## DISCUSSION

Recent advances in scRNA-seq analysis have revealed the cellular and molecular heterogeneity in the human brain organoids (26,27). Comprehensive scRNA-seq analysis of the human cortical brain organoids showed divergent transcriptome profiles between the different protocols (28). Here, we performed comparative scRNA-seq analysis of hBSOs and hMBOs that we previously designed (13,19) and investigated the difference in the transcriptome profiles of each individual cell between hBSOs and hMBOs.

Our comparative analysis suggests that hBSOs and hMBOs contained some unique cell populations. For example, hBSOs preferentially contained INF cluster. The INF cluster expressed genes related to inflammation and immunity, such as *AIF1* and ICAM family (*ICAM-1, ICAM-3*) (Table S2). Microglia cells are the inflammatory and immune cells of the central nervous system, which originate from mesoderm. Microglia, as well as neurons derived from neuroectoderm, are involved in neurodegenerative and neurodevelopmental disorders (29). On the other hand, hMBOs contained MC cluster that expresses genes related to proteoglycan and collagen synthesis, such as BGN and collagen family (*COL1A1, COL1A2*; Table S2). The MC cluster is also detected in human fetal midbrain (Figure 4B). The scRNA-seq analysis of human ESC-derived dopaminergic neurons that were grafted into the rodent brain revealed the presence of fibroblast-like cells (30). These scRNA-seq data are consistent with with our analysis. The two organoids contained different mesodermal-derived cells. The unique transcriptome profile might be related to the different protocols between the two organoids. Further studies using different modalities are required to elucidate the functionality of mesodermal components in the two organoids.

We also compared transcriptome profiles between our two organoids and human fetal midbrain. Our analysis demonstrated that neuronal clusters from hBSOs and hMBOs were highly similar to those of hMB. On the other hand, non-neuronal clusters (e.g. RGC, ENC) from the two organoids showed only moderate similarity to those of hMB (Figure 5B, 5C). The UMAP plot illustrates the two clusters, RGC and OPC, dominantly contained hMB-derived cells (Figure 4B). The RGC cluster expressed genes related to glial cells development, such as *GFAP, FABP7* and *PLP1*. Our protocols are based on current brain organoid protocols for the generation of neuronal cells (31). However, human brain development and neurodegenerative disease are mediated by interactions between neuronal cells and non-neuronal cells, such as glial cells and endothelial cells (29,32,33). In order to expand current research on human brain organoids, it is important to recapitulate not only neuronal cells but also non-neuronal cells.

Since human brain organoids have been established, various protocols for the generation of brain organoids have emerged and have been quickly developed within the decade (31). Region-specific organoids recapitulate the molecular and cellular features of the specific human brain areas (8,9,12,34). In addition to neurons, other cell types such as astrocytes, oligodendrocytes and microglia, are generated in the human brain organoids (5,35,36). The human brain organoid technology has been advanced for the purpose of elucidation of human brain development and brain disorders (4,8). At present, our comparative study mainly focused on the transcriptome profiles of single-cells in hBSOs and hMBOs. We believe that additional comparative analyses on the electrophysiological activities and imaging analysis of the two organoids are needed to evaluate their functionality and to further our understanding human brain development.

## DATA AVAILABILITY

The data that support the findings of this study are available from the corresponding author upon reasonable request.

## SUPPLEMENTAL DATA

Supplementary Data are available at NAR online.

### ACKNOWLEDGEMENT

The authors thank Keren-Happuch E Fan Fen for critical reading of the manuscript.

